# Multisite phosphorylation regulates phenotypic variability in antibiotic tolerance

**DOI:** 10.1101/315085

**Authors:** Elizabeth A. Libby, Shlomi Reuveni, Jonathan Dworkin

## Abstract

Isogenic populations of cells exhibit phenotypic variability that has specific physiological consequences. For example, individual bacteria within a population can differ in their sensitivity to an antibiotic, but whether this variability can be regulated or is generally an unavoidable consequence of stochastic fluctuations is unclear. We observed that a bacterial stress response gene, the (p)ppGpp synthetase *sasA*, exhibits high levels of extrinsic noise in expression, suggestive of a regulatory process. We traced this variability to the convergence of two signaling systems that together control the multisite phosphorylation of a transcription factor, an event largely unexplored in bacteria, This regulatory intersection between a Ser/Thr kinase and a prototypical two component system is crucial for controlling the appearance of outliers, rare cells with unusually high levels of *sasA* expression. Additionally, by examining the full distributions of gene expression we calculated the contribution of the additional Ser/Thr kinase-dependent phosphorylation in setting the relative abundance of cells with a given a level of SasA. We then created a predictive model for the probability of a given cell surviving antibiotic treatment as a function of *sasA* expression. Therefore, our data show that multisite phosphorylation can be used to strongly regulate variations in phenotypes across a bacterial population.

## Introduction

Many bacterial phenotypes, including antibiotic tolerance and virulence, often reflect the phenotype of a subset of the population rather than the average behavior ^1, 2^. Subpopulations of bacteria can arise through purely stochastic processes as well as by regulatory and signaling pathways ^3^. Theoretically, one way to create phenotypic diversity via a signaling pathway is multisite phosphorylation, in which each successive phosphorylation changes the activity of a protein ^4, 5^. However, it has not been experimentally shown in bacterial populations that multisite phosphorylation regulates variation in gene expression between cells, and subsequently, the emergence of phenotypic diversity. Recently, multisite phosphorylation of transcription factors have been observed in pathways involved in antibiotic tolerance and virulence ^6^, suggesting that dynamics of multisite phosphorylation could have particular physiological relevance.

Bacterial signaling is often characterized in the context of two-component signal transduction systems (TCS) that generally consist of a histidine kinase that phosphorylates a response regulator on a single residue, which then acts as a transcription factor ^7^. The stimulus-dependent response of this type of signaling system architecture has been analyzed theoretically ^8, 9^ and experimentally ^10, 11^, with little cell-to-cell variability observed (as quantified by CV), regardless of inducer level. This suggests that extensive cell-to-cell variability is not a general feature of bacterial TCS. However, some notable exceptions have been found for two-component systems with more complex architectures, such as the broad distribution of gene expression in the *E. coli* TorS/TorR regulon ^12^ which has recently been shown to be an important factor for cell survival during oxygen depletion ^13^. The network architecture of bacterial signal transduction systems may therefore play an underappreciated role in the dynamics and survival of bacterial populations.

In addition to TCS, bacteria also have eukaryotic-like (also called Hank’s type) Ser/Thr kinase – phosphatase pairs with homology to eukaryotic kinase systems that perform reversible phosphorylation on Ser and Thr residues ^14^. One particular subfamily of these systems appears to be universally conserved across Gram-positive bacteria and plays key roles in growth and virulence for many clinically important pathogens including the streptococci, *S. aureus*, *M. tuberculosis*, *E. faecalis*, and others ^6, 15^. Genetic and proteomic studies indicate that these Ser/Thr kinases can perform transcriptional regulation of key cellular processes involved in antibiotic tolerance and persistence through multisite phosphorylation of transcription factors. However, to date, the consequences of multisite phosphorylation for gene regulation at the single-cell-level has not been quantified. In this context the model gram-positive bacterium *B. subtilis* presents a comparatively straightforward system to quantify the contribution of the additional Ser/Thr phosphorylation *in vivo*: the homologous kinase-phosphatase pair is PrkC/PrpC, and it has been verified to regulate gene expression through additional phosphorylation of a response regulator.

It has been apparent for over 60 years that bacterial populations contain rare cells that display increased phenotypic resistance to antibiotics ^16^. These cells, presumed to be quiescent, have been implicated in antibiotic treatment failure in genetically susceptible bacterial infections ^17^. To date, it remains unclear to what extent the appearance of these rare cells, is subject to regulation. Emerging evidence strongly implicates elevated levels of the nucleotide second messenger (p)ppGpp as a causative agent of quiescence in many bacterial species ^18-21^. (p)ppGpp downregulates essential cellular processes such as transcription, translation, and DNA replication ^22^. Although the precise mechanism of (p)ppGpp synthesis and its direct cellular targets vary between bacterial species, highly elevated levels of (p)ppGpp confer a quiescent state to the bacterial cell. As many antibiotics target active cellular processes, the resulting cells exhibit increased antibiotic tolerance, suggesting that cell-to-cell variability in (p)ppGpp may be involved in phenotypic resistance to antibiotics ^21, 23-25^.

The mechanistic origin of cell-to-cell variability in (p)ppGpp levels across bacterial populations remains a major open question. To date, this has been best studied in *E. coli*, which has the RelA (p)ppGpp synthetase and the SpoT hydrolase ^26^. In contrast, other bacterial species often possess dedicated (p)ppGpp synthetases, termed small alarmone synthetases (SAS), in addition to bi-functional synthetase-hydrolases ^27^. These SAS proteins can be activated transcriptionally ^22^, suggesting that cell-to-cell variability in (p)ppGpp levels could originate in the transcriptional regulation of the synthetases themselves. In the Gram-positive bacterium *B. subtilis*, (p)ppGpp synthesis is regulated by three distinct proteins: RelA, SasA, and SasB ^28^. *B. subtilis* RelA is a bi-functional (p)ppGpp synthetase-hydrolase, and both SasA and SasB are dedicated synthetases. Although *relA* and *sasB* transcripts are both readily detectable during log phase growth, *sasA* (formerly *ywaC*) transcripts are found at considerably lower levels. However, *sasA* is inducible by certain classes of cell-wall-active antibiotics ^29, 30^, and its induction by alkaline shock increases the cellular levels of ppGpp ^28^. Since *sasA* expression stops growth ^31^, SasA-mediated (p)ppGpp synthesis provides a mechanism to induce cellular quiescence in response to environmental stresses. To date, SasA is only known to be regulated transcriptionally, so significant cell-to-cell variability in *sasA* expression could produce physiologically relevant cell-to-cell variability in (p)ppGpp levels. The pre-existing distribution of *sasA* expression may therefore be critical in predicting the relative survival of cells under conditions that do not specifically induce *sasA*.

In this work, we demonstrate that *sasA* expression displays physiologically relevant amounts of extrinsic noise, although the average level of *sasA* expression is very low during growth under non-inducing conditions. Furthermore, we find that both the distribution of *sasA* expression and the frequency of outliers are strongly regulated by the activity of a highly conserved eukaryotic-like Ser/Thr kinase system and its subsequent multisite phosphorylation of a transcription factor. Using quantitative analysis of the full distributions of *sasA* expression, we find that multisite phosphorylation is responsible for exponentially regulating the abundance of cells with a given level of SasA and generate a predictive model for *sasA*-expression-dependent antibiotic tolerance.

## Results

### The (p)ppGpp synthetase *sasA* exhibits high levels of extrinsic noise in expression

While the population average level of *sasA* expression during growth is extremely low ^29^, the average behavior may mask important phenotypic variation between cells. We therefore generated a transcriptional reporter for *sasA* (P_*sasA*_-*yfp*) to study the population at the single-cell level. Surprisingly, there was considerable cell-to-cell variability in P_*sasA*_-*yfp* (coefficient of variation, CV~4.95 ± 0.42, mean ± SEM), with most cells having very low expression, and rare cells showing significantly higher levels of expression (Fig. 1A). Quantification of YFP fluorescence revealed that a small fraction of the population had much higher (>~10x) levels of fluorescence than the mean, and rare cells had ~100x. Note that a typical bacterial gene has a CV in the range 0.1-1 ^32-34^.

**Figure 1:**
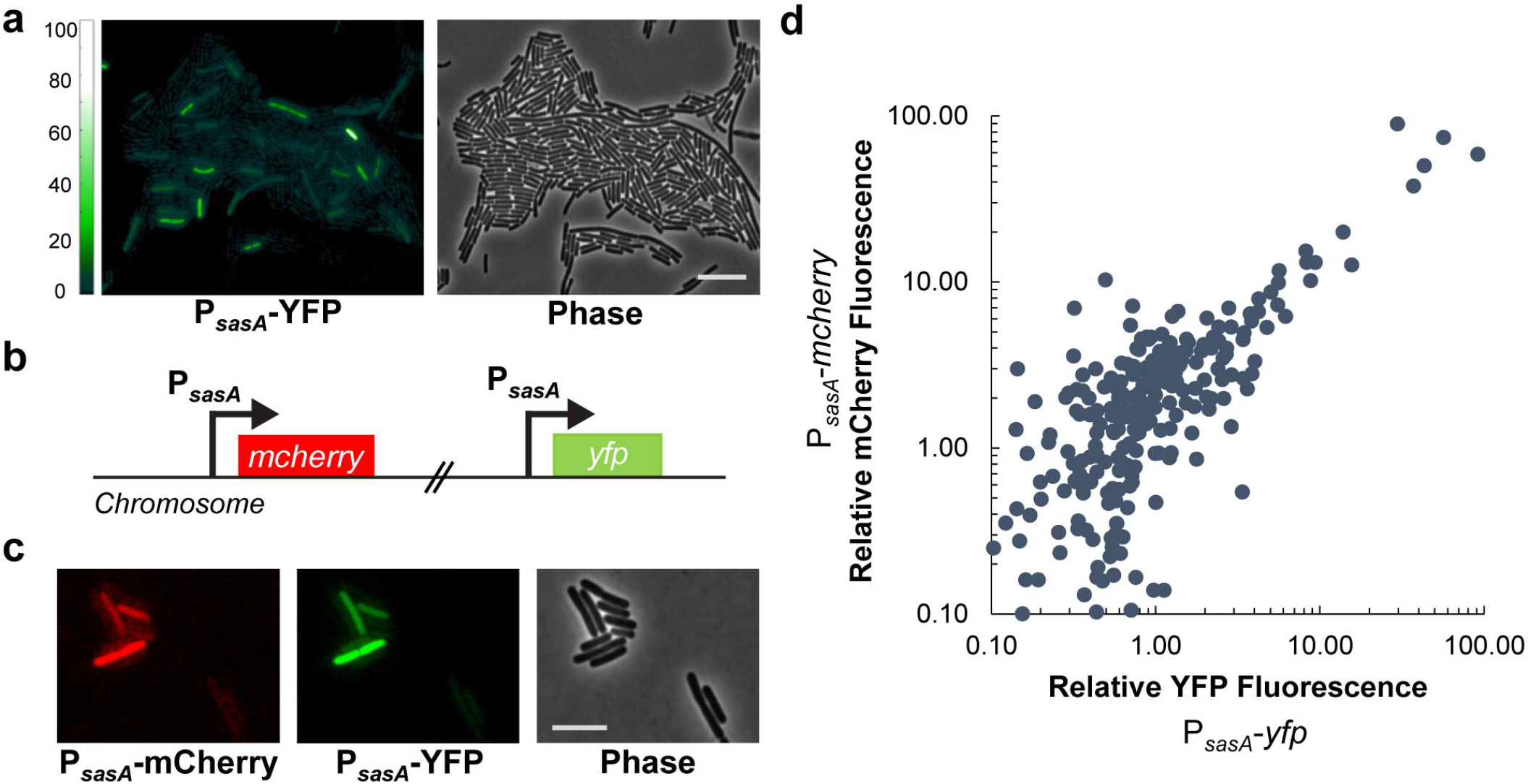
*sasA* exhibits cell-to-cell variability in expression. **A)** P_*sasA*_-*yfp* expression in log phase culture. **Left:** YFP image. Color scale indicates the fold change relative to the population average. **Right:** Phase contrast. Scale bar indicates 10 μm. **B)** Schematic of dual-color transcriptional reporter strain to characterize the noise in *sasA* expression. Two copies of the *sasA* promoter driving different reporters (P_*sasA*_-*yfp*, P_*sasA*_-*mcherry*) are inserted at separate ectopic loci. **C)** Evidence for extrinsic noise in *sasA* expression. Images of single cells from the dual reporter strain with significant fluorescence shown in mCherry (**Left**), YFP (**Center**), and phase contrast (**Right**) channels. Scale bar indicates 5 μm. **D)** Quantification of cellular fluorescence intensities for single cells in the experiment described in **B, C**. Values were normalized relative to the population mean value for each channel. Data shown was measured on ~ 580 cells with detectable fluorescence in both channels. The fluorescence of the two reporters have a Pearson’s correlation coefficient of r ~ 0.90±0.08 (mean ± SEM, 4 experiments). For each reporter, the CV ~ 4.95±0.42 (mean ± SEM, 4 experiments).

The high levels of cell-to-cell variability in *sasA* expression could be caused by intrinsic noise from the promoter itself, or by extrinsic noise originating in an upstream process ^35^. To differentiate between these mechanisms, we used a strain with dual fluorescent reporters, P_*sasA*_-*yfp* and P_*sasA*_-*mcherry* (Fig. 1B). Expression of the dual reporters in individual cells was highly correlated (Pearson’s correlation coefficient, r~0.90±0.08, mean ± SEM), demonstrating that the noise was largely extrinsic to the promoter (Fig. 1C, D).

To determine if variation in YFP levels were simply caused by global changes in the population that result in the accumulation of large amounts of fluorescent protein ^36^, P_*sasA*_-*yfp* was compared to a presumably unrelated promoter known to be constitutively active during log phase growth, P_*veg*_-*mcherry* ^37^ (Fig. S1). We found that YFP and mCherry levels were not highly correlated, suggesting that the high levels of variability in *sasA* expression are largely caused by a *sasA*-specific pathway. We then tested a previously characterized *sasA* regulator, the sigma factor σ^M^ (SigM) ^38^, that is required for *sasA* expression ^30^ (Fig. S2A, B). However, *sasA* expression also did not correlate strongly with *sigM* expression (Fig. S2C, D) demonstrating that *sigM* levels alone do not predict variability in *sasA* expression.

### *sasA* is repressed by the Ser/Thr kinase PrkC through the response regulator WalR

Another potential regulator of *sasA* is the WalR transcription factor observed to bind the *sasA* promoter in a genome-wide screen ^39^. WalR is the response regulator of the essential WalRK two-component system and is activated by phosphorylation of Asp-53 by WalK ^40^. Once phosphorylated, WalR can either activate or repress genes in its regulon. A reversible second phosphorylation on WalR Thr-101 by the eukaryotic-like Ser/Thr kinase-phosphatase pair PrkC/PrpC ^41^ further increases WalR activity at both activating and repressing sites ^42^. In rich media (LB), the multisite phosphorylation of WalR affects gene expression (*e.g.*, enhanced activation of *yocH*) specifically in post-log phase ^42^. However, in commonly used defined minimal media (S7-glucose), there is a consistent PrkC-dependent effect on the population average level of *yocH* expression throughout log phase (Fig. S3). *sasA* is known to be activated by antibiotics such as bacitracin ^29^ through σ^M^ activation. We first tested whether the PrkC/PrpC – WalR system regulates *sasA* at the population level to determine if WalR activates or represses *sasA*. We found that PrkC activity represses *sasA* expression through WalR Thr101~P during bacitracin treatment (Fig. S4). Based on these bulk measurements, we developed a model for *sasA* regulation (Fig. 2A) in which PrkC activity further potentiates WalR-repressing activity at *sasA* through a second phosphorylation of WalR at Thr-101. However, it remained unclear to what extent multisite phosphorylation of WalR affects pre-existing cell-to-cell variability in *sasA* expression under non-inducing conditions.

**Figure 2:**
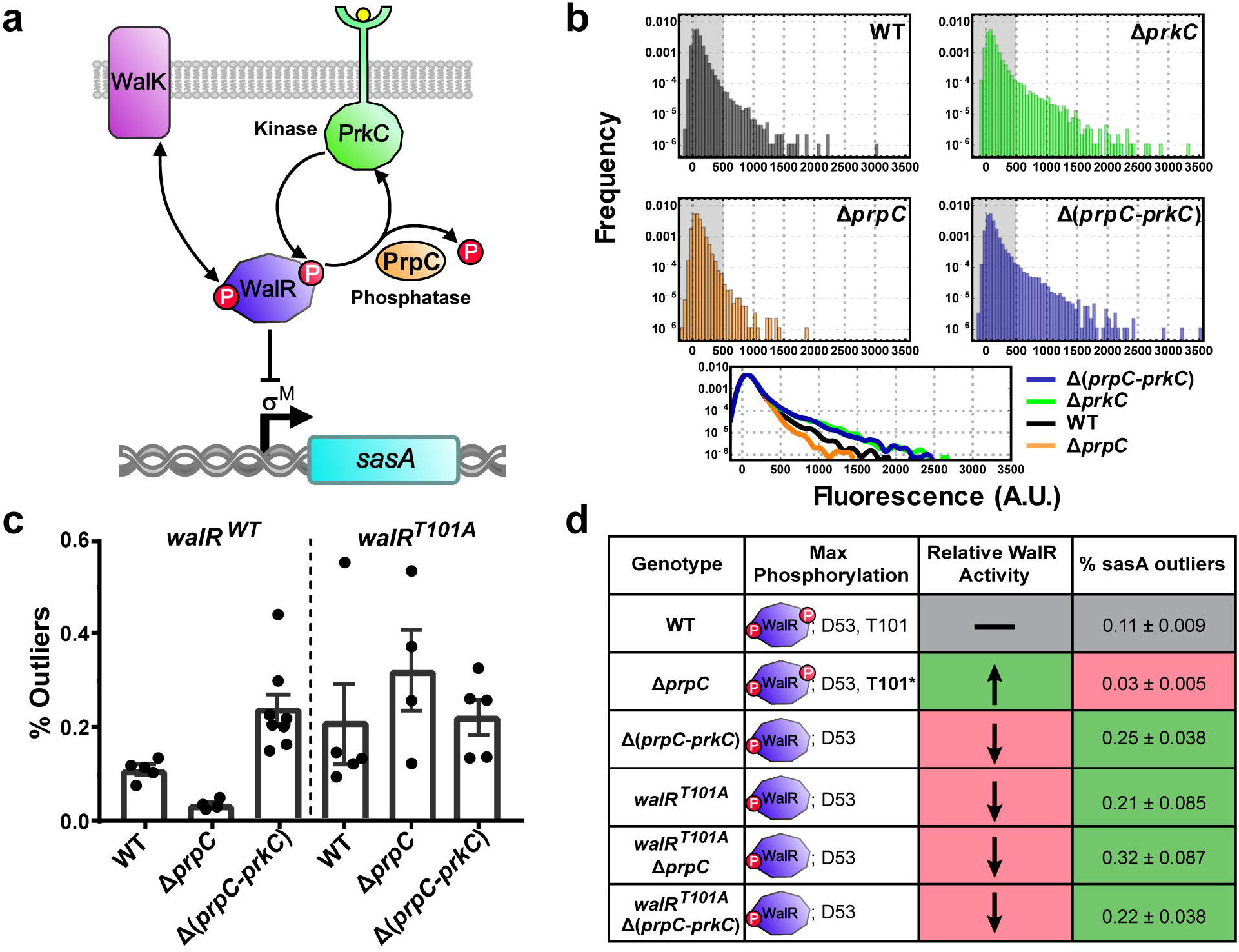
The Ser/Thr kinase PrkC and phosphatase PrpC regulate cell-to-cell variability in *sasA*. **A)** Model for PrkC-dependent regulation of *sasA*. WalR binding to the *sasA* promoter represses *sasA*. WalR activity is primarily controlled by phosphorylation on Asp-53 by its cognate histidine kinase WalK, and secondarily by phosphorylation on Thr-101 by the Ser/Thr kinase PrkC. WalR Thr-101~P can be dephosphorylated by the phosphatase PrpC. Thr-101~P further enhances the repressing activity of WalR Asp-53~P, resulting in lower expression of *sasA* under conditions with high Thr-101~P (e.g., Δ*prpC*). Under conditions lacking Thr-101~P (*e.g.*, Δ*prkC*), the increased repression of *sasA* is relieved. **B)** Histograms demonstrating PrkC-dependent cell-to-cell variability in *sasA*. **Top:** P_*sasA*_-*yfp* reporter activity was quantified by flow cytometry in wild type (WT, gray), Δ*prpC* (orange), Δ*prkC* (green), and Δ(*prpC-prkC*) (blue) backgrounds. Shaded region of each plot indicates the range of cellular autofluorescence observed. Each histogram was computed from data on ~3.0*10^4^ events. **Bottom:** Smoothed, overlaid histograms of the data shown above for comparison. **C)** Percentage of outliers in each genetic background. At least 4 independent experiments, similar to and including the representative one shown in **B**, were performed in *walR ^WT^* (**Left**) and *walR ^T101A^* (**Right**) backgrounds. Each experiment was normalized to a control and outliers were defined as cells above a fixed threshold level of normalized fluorescence (~1250 A.U.). Dots represent the percentage of each population that is above the threshold; bars and lines represent the mean and SEM, respectively. **D)** Summary of each genotype, its effect on the maximal occupancy of the two known WalR phosphosites, and the expected effect on WalR activity relative to WT. For each genotype the mean percentage of outliers in *sasA* expression (shown in **C**) are also summarized. Note that in the Δ*prpC* background, the Thr-101 phosphorylation is stabilized (denoted as **T101\***).

### PrkC regulates noise in *sasA* expression through WalR Thr-101 phosphorylation

Cell-to-cell variability in gene expression can be tuned by changing repressor-binding affinities ^43, 44^, suggesting that multisite phosphorylation of WalR may play a critical role in setting the observed distribution of *sasA* expression across the population. To test this, we measured the distribution of *sasA* expression in wild type (WT) cells and compared it to genetic backgrounds that alter the phosphorylation state of WalR: Δ*prpC* (no phosphatase, high levels of T101~P), and Δ*prkC* (no kinase, no detectable T101~P) (Fig. 2B). Qualitatively, in the Δ*prpC* background, the frequency of cells with high levels of *sasA* expression was strongly reduced, whereas it was strongly increased in the Δ*prkC* background. The PrpC-dependent effect on *sasA* expression requires PrkC, since the distribution of *sasA* expression in a strain lacking both the kinase and phosphatase (Δ(*prpC-prkC*)) is very similar to a strain lacking just the kinase.

We first sought to quantify the effect of WalR multisite phosphorylation on the frequency of “outliers”: cells with a level of *sasA* expression above a fixed threshold in each population. We therefore compared independent measurements of the distribution of *sasA* expression in WT, Δ*prpC*, and Δ(*prpC-prkC*) backgrounds (Fig. 2C, left) and found that PrkC significantly affects the mean frequency of outliers >8 fold by this measure (*walR ^WT^*, Δ*prpC* vs Δ(*prpC-prkC*): **p-value~0.004, Kolmogorov-Smirnov test). We repeated the measurements in a *walR ^T101A^* background (Fig. 2C, right) and found that PrkC no longer has a significant effect on the mean frequency of outliers in the phosphosite mutant background (*walR ^T101A^*, Δ*prpC* vs Δ(*prpC-prkC*): p-value~0.56, ns, Kolmogorov-Smirnov test). These results are consistent with increased WalR activity by Thr-101 phosphorylation causing increased repression of *sasA*, and thereby regulating the frequency of *sasA* outliers (Fig. 2D). Furthermore, heterologous expression of PrkC was sufficient to reduce the frequency of outliers observed in the Δ*prkC* background (Fig. S5A). Heterologous expression of PrkC was also able to further reduce the variability to below that observed in the Δ*prpC* background, approaching the level of cellular autofluorescence (Fig. S5B). This suggests that at least some of the remaining variability in the Δ*prpC* background arises due to incomplete saturation of WalR T101~P.

This definition of outliers, however, relies on the definition of a cutoff threshold and therefore does not fully address how multisite phosphorylation affects the entire distribution of *sasA* expression across the population. To quantify the effect of PrkC on the distribution of cell-to-cell variability in *sasA* expression, we deconvolved the measured data from the cellular autofluorescence (SI). This resulted in autofluorescence-free distributions of *sasA* expression, allowing better quantitative comparison of expression between genetic backgrounds (Fig. 3A). This deconvolution method uses only the first two moments (*i.e*., the mean and variance) of the observed distributions of fluorescence. As such, the auto-fluorescence free distributions are relatively insensitive to the observed “outliers” in each distribution, but makes a statistical prediction for those frequencies. To verify the accuracy of the predictions, these calculated distributions were re-convolved with the cellular autofluorescence and the reconstructed data set compared to the original data (Fig. S6). Calculation of the relative enrichment of cells with a given level of *sasA* in each genetic background revealed that maximal T101~P (Δ*prpC*) compared to the absence of T101~P (Δ*prkC*) results in exponential changes in the relative abundance of cells at a given level of *sasA* (Fig. 3B).

**Figure 3:**
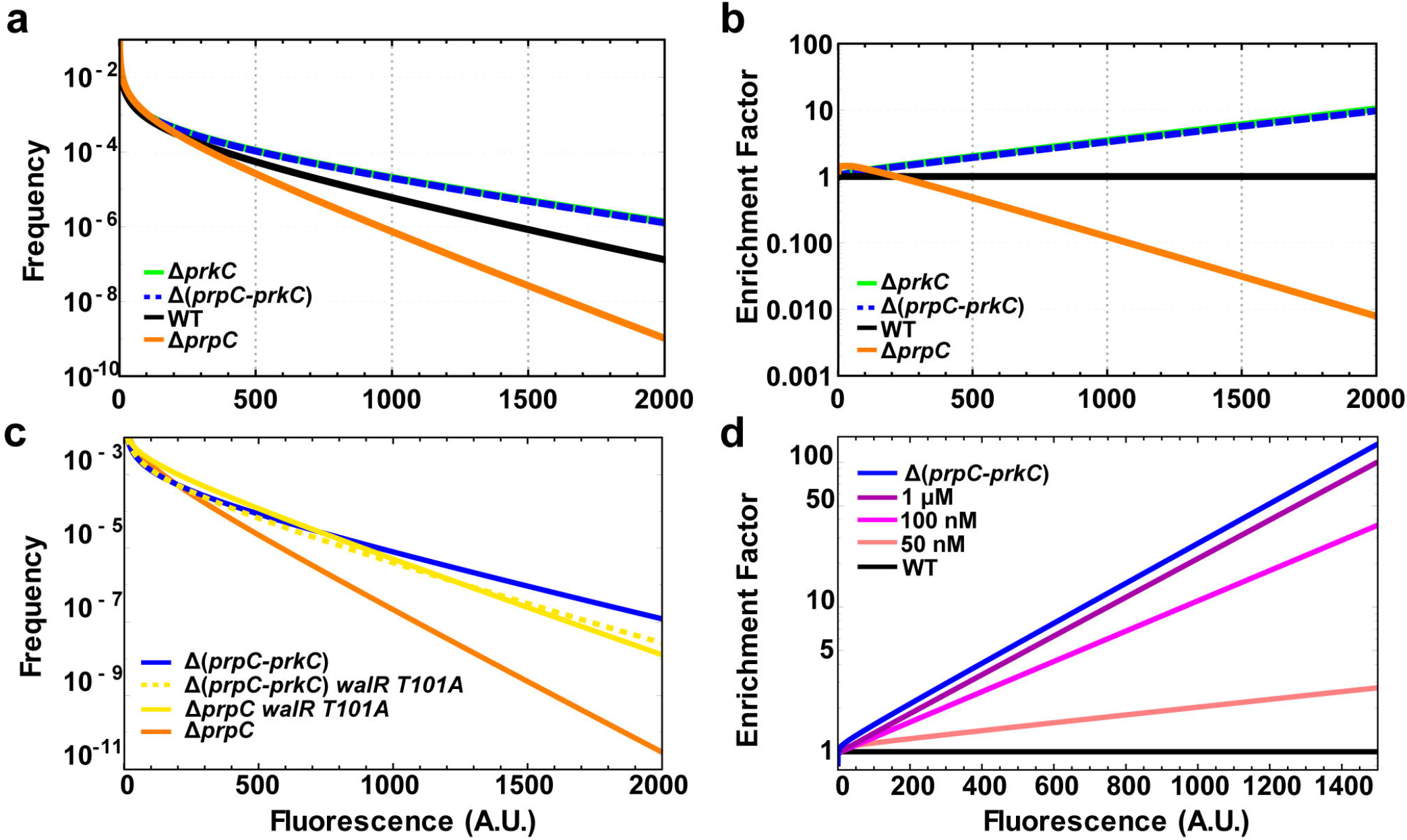
Multisite phosphorylation causes exponential changes in abundance of cells with a given level of *sasA* expression. **A)** Functional fits of autofluorescence-free distributions of *sasA* expression. A deconvolution algorithm was used to remove the contribution of autofluorescence from the measured distributions of *sasA* expression shown in Figure 2B. Validation of the fits are shown in Fig. S6. **B)** Effect of PrkC on the distribution of *sasA* expression. For each genotype in **A** the frequency of cells with a given level of *sasA* expression was normalized against the equivalent frequency in the WT population. On this semi-logarithmic plot, straight lines indicate that the enrichment factor changes ~exponentially with fluorescence. **C)** PrkC-dependent regulation of cell-to-cell variability in *sasA* is WalR Thr-101-dependent. P_*sasA*_-*yfp* reporter activity was quantified by flow cytometry in a Δ*prpC walR T101A* (yellow, solid) background and compared to Δ*prpC* (orange), Δ(*prpC-prkC*) (blue), and Δ(*prpC-prkC*) *walR T101A* (yellow, dashed) backgrounds in the same experiment. Plots were generated by applying the deconvolution method described in **A** to the measured data in Fig. S7A. **D)** Gradual inhibition of PrkC results in progressive enrichment of cells with elevated levels of *sasA*. P_*sasA*_-*yfp* reporter activity was quantified by flow cytometry during treatment with increasing concentrations of staurosporine: 0 (solvent only; black), 50 nM, 100 nM, and 1 μM (shades of magenta), in otherwise WT populations. For reference, a Δ(*prpC-prkC*) (blue) population treated with solvent only is shown; note that the effect of staurosporine saturates at this level. Plots were generated by applying the deconvolution method described in **A** to the measured data in Fig. S7B, C. For each condition the frequency of cells with a given level of *sasA* expression was normalized against the equivalent frequency in the WT population. As in **B**, straight lines indicate that enrichment increases ~exponentially with fluorescence. Enrichment becomes more pronounced as the concentration of staurosporine increases and saturates at the level of complete inhibition ~Δ(*prpC-prkC*).

Together, the model and the outlier analysis in Fig. 2 suggest that PrkC-dependent regulation of the distribution of *sasA* expression requires the second WalR phosphosite at Thr-101. To test this, we repeated the deconvolution procedure for a strain expressing a WalR mutant that lacks the Thr-101 phosphosite, *walR T101A*, and found that the PrkC-dependent effect on *sasA* expression is indeed WalR Thr-101-dependent (Fig. 3C, Fig. S7A). Thus, multisite phosphorylation is responsible for the exponential depletion of cells with medium to high levels of *sasA* expression in the Δ*prpC* background (Fig. 3B).

We then measured how intermediate levels of multisite phosphorylation regulate the distribution of *sasA* expression using the kinase inhibitor staurosporine to progressively inhibit PrkC activity ^45^ (Figs. S7B; S8). The distributions of *sasA* expression were again deconvolved (Fig. S7C), and we calculated the relative enrichment of cells with a given level of *sasA* fluorescence at increasing concentrations of staurosporine (Fig. 3D). Titration of PrkC activity resulted in exponential enrichment of cells with a given level of *sasA*. Therefore, even small changes in PrkC activity result in large changes in the abundance of “outliers”, cells with unusually high levels of *sasA*.

### *sasA* expression level continuously predicts the probability of surviving antibiotic treatment

Cell-to-cell variability in (p)ppGpp production has been proposed to result in cell-to-cell variability in antibiotic survival ^47,48, 49^. However, a direct and quantitative relationship between the expression of a transcriptionally regulated (p)ppGpp synthetase and the probability of survival for an individual cell has not been demonstrated. We therefore sought to determine if cells with pre-existing high levels of *sasA* preferentially survive antibiotic exposure, and if so, provide a model for how the level of *sasA* expression influences the probability of survival for a given cell.

We used ciprofloxacin, a DNA gyrase inhibitor that does not significantly increase the population average level of *sasA* expression ^29^. We measured (Fig. S9) and deconvolved (Fig. 4A, B) the distributions of *sasA* expression (P_*sasA*_-*yfp*) both pre-and post-ciprofloxacin treatment that results in ~99% killing in both WT and Δ*sasA* backgrounds. We note that, importantly, the starting distributions of P_*sasA*_-*yfp* are very similar in both genetic backgrounds, allowing a direct comparison. Using these distributions, we calculated the relative enrichment of cells with a given level of *sasA* following antibiotic treatment (Fig. 4C), yielding a simple model for the effect of antibiotic treatment on the distribution of *sasA* expression (SI). Because survival after antibiotic treatments can be affected by many processes, we separated out the size of the *sasA*-dependent effect by using a Δ*sasA* mutant as a control. WT populations exhibited a significant increase in the fraction of cells with elevated levels of *sasA* (Fig. 4C). This effect is strongly reduced in the Δ*sasA* background, demonstrating that *sasA* has a significant contribution to survival after ciprofloxacin treatment.

**Figure 4:**
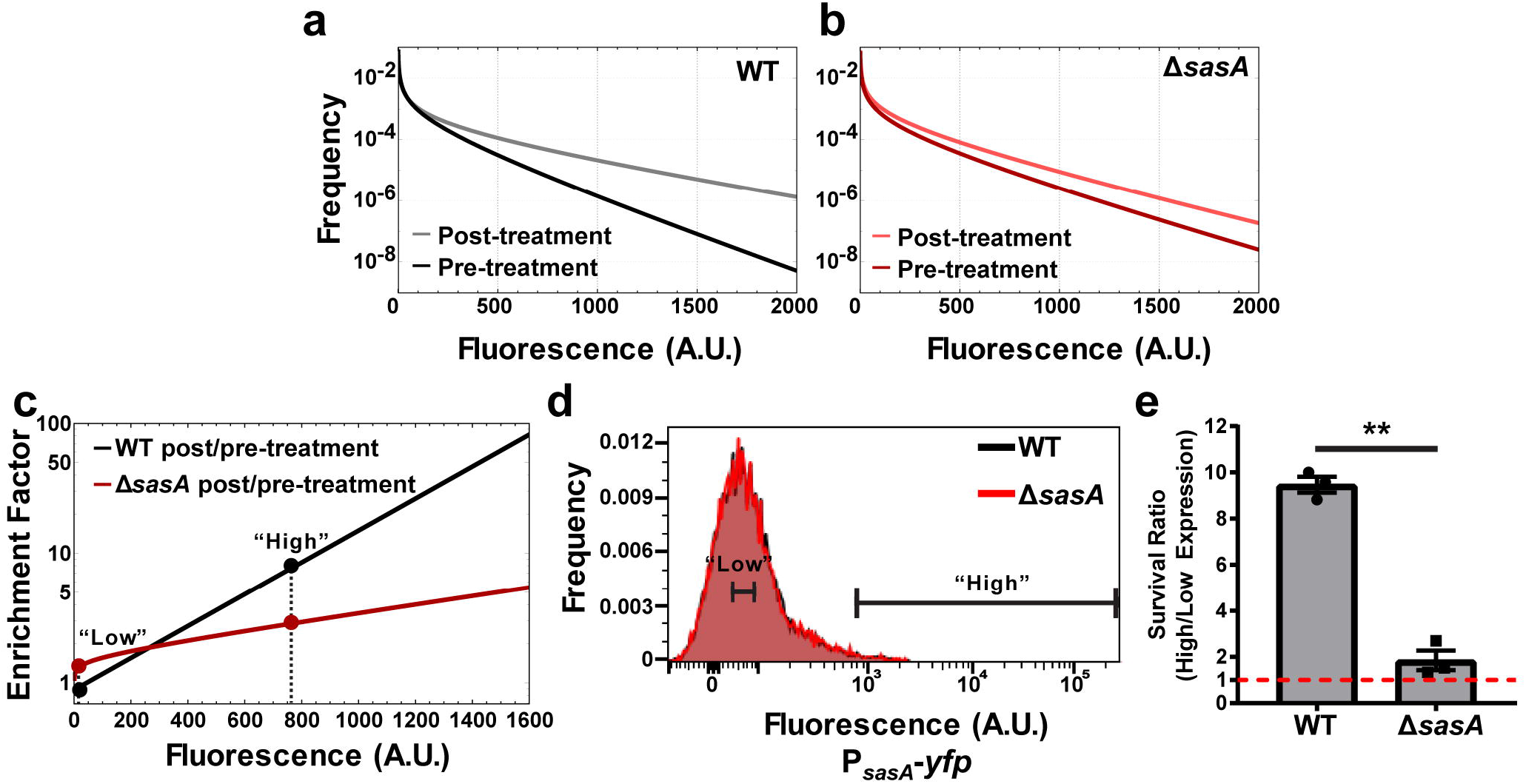
Cells with elevated *sasA* expression preferentially survive antibiotic treatment. **A)** Autofluorescence-free distributions of *sasA* expression before (dark gray) and after (light gray) ciprofloxacin treatment in an otherwise WT background based on the data shown in Fig. S9A. **B)** Autofluorescence-free distributions of *sasA* expression before (dark red) and after (light red) ciprofloxacin treatment in a Δ*sasA* background based on the data shown in Fig. S9B. **C)** Enrichment plot following antibiotic treatment for WT and Δ*sasA* populations. For each genotype in **A** and **B** the frequency of cells with a given level of P_*sasA*_-*yfp* after antibiotic treatment was normalized against the corresponding frequency prior to antibiotic treatment. Dashed lines and dots indicate the average values used in **D** and **E**. **D)** Histograms of P_*sasA*_-*yfp* fluorescence in WT (black) and Δ*sasA* (red) backgrounds measured by flow cytometry prior to sorting into “low” (4.1 ± 1.8) or “high” (779.5 ± 134.9) expression groups (relative mean ± SEM, 3 experiments). Each histogram is comprised of data obtained from ~3.0*10^4^ events and is from a representative experiment used in **E**. **E)** Relative survival of the “high” and “low” expression populations in **D** after treatment with ciprofloxacin for ~3.5 h as in **A, B**. Bars indicate the mean survival ratio of the “high” to “low” expressing populations. The data shown represents 3 independent experiments (paired t-test, p-value ~ 0.009, **). The red dashed line represents a survival ratio of 1, indicating no advantage.

We then sought to determine whether our enrichment model reflects the probability of survival for cells as a function of *sasA* expression. Since SasA is a (p)ppGpp synthetase, the *sasA*-dependent enrichment we observe post-ciprofloxacin treatment could be due to an expression-dependent probability of surviving antibiotic treatment. Alternately, since the measured increase in mean fluorescence was relatively small (~2 fold), it is also possible that ciprofloxacin acts in a complex, expression-dependent, manner to generate the observed post-treatment distribution of *sasA* expression but does not affect survival. To differentiate between these hypotheses, we used FACS to sort bacteria prior to ciprofloxacin treatment from both WT and Δ*sasA* populations into “high” (upper ~1%) and “low” (~average) P_*sasA*_-*yfp* expression groups (Fig. 4D). Following ciprofloxacin treatment (as in Figs. 4A, B, S9), the relative survival of “high” and “low” expression cells, or the survival ratio, was assayed by plating for CFUs (Fig. 4E). The average fluorescence cutoff values used in the FACS experiments, low: 4.1±1.8 and high: 779.5±134.9 (mean ± SEM, 3 experiments), were then used as inputs for a model where the enrichment of cells with increased levels of *sasA* (Fig. 4C) is caused by increased survival (SI). The model yielded good agreement with the results of the FACS experiments: it predicted relative survival ratios of ~9 for wild-type, and ~2 for Δ*sasA*, respectively, compared to the measured values of ~9.5 ± 0.6 (WT) and 1.8 ± 0.7 (Δ*sasA*) (Fig. 4E). Therefore, the enrichment of cells with elevated levels of *sasA* post-ciprofloxacin treatment can be largely attributed to the increased survival probability of pre-existing cells in the population with elevated *sasA* expression.

Taken together, our results demonstrate that an important consequence of PrkC-dependent multisite phosphorylation of WalR is the regulation of cell-to-cell variability, or noise, in the WalR regulon gene *sasA*. By comparing the full distributions of gene expression, we demonstrate that this effect is not just confined to the regulation of outliers in gene expression above an arbitrary threshold within the population, but has an exponential effect on the relative abundance of cells with a given level of expression. By analyzing the full distributions of expression, we are also able to demonstrate that *sasA* expression also continuously affects the antibiotic tolerance of individual cells: specifically, the survival probability during a fixed course antibiotic treatment. This model (SI) is consistent with cell sorting experiments that explicitly demonstrate that the observed distributions are a consequence of survival probability.

## Discussion

Antibiotic tolerance is believed to be an important factor in the failure of antibiotic treatments and a key step toward the development of antibiotic resistance ^50^. We therefore sought to trace the origin of the cell-to-cell variability in expression of *sasA* and determine if it can be regulated by genetic or chemical means. Noise in gene expression can be conceptually separated into intrinsic and extrinsic noise. Although it is difficult to design strategies to specifically target events generated by intrinsic noise, extrinsic noise may have upstream regulatory pathways that can be modulated. Therefore, it is significant that the cell-to-cell variability in *sasA* was dominated by extrinsic noise at high levels of expression (Fig. 1) that have the strongest effect on antibiotic tolerance (Fig. 4). Furthermore, since multisite phosphorylation is responsible for setting the observed distribution of cell-to-cell variability (Figs. 2, 3), this regulatory pathway could be a novel antibiotic target.

Multisite phosphorylation can expand the range of a protein’s function, generating both switch-like ^51, 52^, and graded ^53, 54^ changes in average activity. In contrast, here we observed only minimal changes in the average levels of *sasA* expression as a function of PrkC activity, but measured up to a ~100-fold effect on the frequency of “outliers”, cells with particularly high levels of expression (Fig. 3B). This response was shown to be graded, rather than switch-like, likely arising as a consequence of the integration of signals from two distinct signaling systems. A single phosphorylation at WalR Asp-53 strongly, but imperfectly, represses the *sasA* promoter. The addition of the second phosphorylation at Thr-101 by a distinct signaling system then acts as a second input to further regulate WalR. Interestingly, even small changes in activity of the second system result in marked changes in the frequency of outliers. Heterologous expression of PrkC is capable of reducing the variability observed to close to cellular autofluorescence, but does not eliminate it completely. This remaining variability in *sasA* may be due to PrkC overexpression still being unable to completely saturate WalR phosphorylation, intrinsic noise at the promoter, or as yet unidentified sources.

Transcriptional regulation of outliers in eukaryotes has been shown to be predictive of which cancer cells survive drug treatment ^55^. Here we found that transcriptional regulation by multisite phosphorylation is also critical for setting the pre-existing distribution of survival probabilities for cells within a bacterial population. Distinct from bacterial persistence, which is characterized by bi-phasic killing, these survival probabilities reflect antibiotic tolerance or the killing kinetics during a relatively short, fixed time-course, antibiotic treatment ^56^. In the Δ*sasA* background, we observed a much weaker dependence of antibiotic tolerance on *sasA* expression. This residual dependence is consistent with previous results that have implicated many global processes in antibiotic tolerance including heterogeneity in growth state ^24, 57, 58^ and enhanced expression of drug efflux pumps ^59, 60^. This is also consistent with the relatively weak correlation in expression between *sasA* and the constitutive promoter *veg* (Fig. S2). Indeed, it remains to be seen precisely how cellular physiology changes in a *sasA*-expression dependent manner. SasA has been shown to be important for ribosome assembly in *B. subtilis* ^31^ and for survival during envelope stress in *S. aureus* ^61^. More generally, various cellular processes are known to be directly and indirectly affected by rising (p)ppGpp levels including inhibition of DNA primase activity ^62^, and reduction in intracellular GTP pools ^63^ thereby downregulating rRNA transcription ^64^. As our results show that antibiotic survival increases continuously with *sasA* expression, it suggests that SasA exerts a continuous effect proportional to its level on physiological processes that mediate ciprofloxacin killing. Therefore, multisite phosphorylation may provide a “bet-hedging” strategy to regulate the phenotypic diversity of a bacterial population, serving as a broadly useful mechanism to tune the frequency of rare phenotypes that facilitate survival under adverse conditions.

## Acknowledgements

We thank Eric Brown for the strain EB1385, Isaac Plant (Silver lab) for pIP384 and IP563, and Amir Figueroa at the Microbiology & Immunology Core Facility at Columbia University for assistance with flow cytometry and cell sorting. This work was supported by NIH GM114213 and a BWF Investigators in the Pathogenesis of Infectious Disease award to JD, a grant from the Department of Systems Biology at Harvard Medical School to EAL, and SR was supported by NIH GM095784 and gratefully acknowledges support from the Azrieli Foundation. Author Contributions: Conceived and designed the experiments EAL and JD. EAL performed the experiments. EAL and SR analyzed data. Contributed reagents/materials/analysis tools: EAL and SR. Wrote the paper: EAL, SR and JD.

## References

1 Arnoldini, M. et al. Bistable expression of virulence genes in salmonella leads to the formation of an antibiotic-tolerant subpopulation. PLoS Biol 12, e1001928, doi:10.1371/journal.pbio.1001928 (2014).

2 Norman, T. M., Lord, N. D., Paulsson, J. & Losick, R. Stochastic Switching of Cell Fate in Microbes. Annual review of microbiology 69, 381–403, doi:10.1146/annurev-micro-091213-112852 (2015).

3 Maamar, H., Raj, A. & Dubnau, D. Noise in gene expression determines cell fate in Bacillus subtilis. Science 317, 526–529, doi:10.1126/science.1140818 (2007).

4 Gunawardena, J. Multisite protein phosphorylation makes a good threshold but can be a poor switch. Proc Natl Acad Sci U S A 102, 14617–14622, doi:10.1073/pnas.0507322102 (2005).

5 Ortega, F., Garces, J. L., Mas, F., Kholodenko, B. N. & Cascante, M. Bistability from double phosphorylation in signal transduction. Kinetic and structural requirements. FEBS J 273, 3915–3926, doi:10.1111/j.1742-4658.2006.05394.x (2006).

6 Wright, D. P. & Ulijasz, A. T. Regulation of transcription by eukaryotic-like serine-threonine kinases and phosphatases in Gram-positive bacterial pathogens. Virulence 5, 863–885, doi:10.4161/21505594.2014.983404 (2014).

7 Laub, M. T. & Goulian, M. Specificity in two-component signal transduction pathways. Annu Rev Genet 41, 121–145, doi:10.1146/annurev.genet.41.042007.170548 (2007).

8 Batchelor, E. & Goulian, M. Robustness and the cycle of phosphorylation and dephosphorylation in a two-component regulatory system. Proc Natl Acad Sci U S A 100, 691–696, doi:10.1073/pnas.0234782100 (2003).

9 Shinar, G., Milo, R., Martinez, M. R. & Alon, U. Input output robustness in simple bacterial signaling systems. Proc Natl Acad Sci U S A 104, 19931–19935, doi:10.1073/pnas.0706792104 (2007).

10 Batchelor, E., Silhavy, T. J. & Goulian, M. Continuous control in bacterial regulatory circuits. J Bacteriol 186, 7618–7625, doi:10.1128/JB.186.22.7618-7625.2004 (2004).

11 Miyashiro, T. & Goulian, M. High stimulus unmasks positive feedback in an autoregulated bacterial signaling circuit. Proc Natl Acad Sci U S A 105, 17457–17462, doi:10.1073/pnas.0807278105 (2008).

12 Roggiani, M. & Goulian, M. Oxygen-Dependent Cell-to-Cell Variability in the Output of the Escherichia coli Tor Phosphorelay. J Bacteriol 197, 1976–1987, doi:10.1128/JB.00074-15 (2015).

13 Carey, J. N. et al. Regulated Stochasticity in a Bacterial Signaling Network Permits Tolerance to a Rapid Environmental Change. Cell, doi:10.1016/j.cell.2018.02.005 (2018).

14 Pereira, S. F., Goss, L. & Dworkin, J. Eukaryote-like serine/threonine kinases and phosphatases in bacteria. Microbiol Mol Biol Rev 75, 192–212, doi:10.1128/MMBR.00042-10 (2011).

15 Manuse, S., Fleurie, A., Zucchini, L., Lesterlin, C. & Grangeasse, C. Role of eukaryotic-like serine/threonine kinases in bacterial cell division and morphogenesis. FEMS Microbiol Rev 40, 41–56, doi:10.1093/femsre/fuv041 (2016).

16 Bigger, J. W. Treatment of staphylococcal infections with penicillin by intermittent sterilisation. Lancet 244, 497–500 (1944).

17 Fauvart, M., De Groote, V. N. & Michiels, J. Role of persister cells in chronic infections: clinical relevance and perspectives on anti-persister therapies. J Med Microbiol 60, 699–709, doi:10.1099/jmm.0.030932-0 (2011).

18 Abranches, J. et al. The molecular alarmone (p)ppGpp mediates stress responses, vancomycin tolerance, and virulence in Enterococcus faecalis. J Bacteriol 191, 2248–2256, doi:10.1128/JB.01726-08 (2009).

19 Gaca, A. O. et al. Basal levels of (p)ppGpp in Enterococcus faecalis: the magic beyond the stringent response. MBio 4, e00646–00613, doi:10.1128/mBio.00646-13 (2013).

20 Dordel, J. et al. Novel determinants of antibiotic resistance: identification of mutated loci in highly methicillin-resistant subpopulations of methicillin-resistant Staphylococcus aureus. MBio 5, e01000, doi:10.1128/mBio.01000-13 (2014).

21 Maisonneuve, E. & Gerdes, K. Molecular mechanisms underlying bacterial persisters. Cell 157, 539–548, doi:10.1016/j.cell.2014.02.050 (2014).

22 Liu, K., Bittner, A. N. & Wang, J. D. Diversity in (p)ppGpp metabolism and effectors. Curr Opin Microbiol 24, 72–79, doi:10.1016/j.mib.2015.01.012 (2015).

23 Nguyen, D. et al. Active starvation responses mediate antibiotic tolerance in biofilms and nutrient-limited bacteria. Science 334, 982–986, doi:10.1126/science.1211037 (2011).

24 Balaban, N. Q., Merrin, J., Chait, R., Kowalik, L. & Leibler, S. Bacterial persistence as a phenotypic switch. Science 305, 1622–1625, doi:10.1126/science.1099390 (2004).

25 Maisonneuve, E., Castro-Camargo, M. & Gerdes, K. (p)ppGpp Controls Bacterial Persistence by Stochastic Induction of Toxin-Antitoxin Activity. Cell 154, 1140–1150, doi:10.1016/j.cell.2013.07.048 (2013).

26 Cashel, M., Gentry, D., Hernandez, V. & Vinella, D. in Esherichia coli and Salmonella; Cellular and Molecular Biology Vol. 2 1458–1496 (ASM Press, Washington DC, 1996).

27 Atkinson, G. C., Tenson, T. & Hauryliuk, V. The RelA/SpoT homolog (RSH) superfamily: distribution and functional evolution of ppGpp synthetases and hydrolases across the tree of life. PLoS One 6, e23479, doi:10.1371/journal.pone.0023479 (2011).

28 Nanamiya, H. et al. Identification and functional analysis of novel (p)ppGpp synthetase genes in Bacillus subtilis. Mol Microbiol 67, 291–304, doi:10.1111/j.1365-2958.2007.06018.x (2008).

29 D’Elia, M. A. et al. Probing teichoic acid genetics with bioactive molecules reveals new interactions among diverse processes in bacterial cell wall biogenesis. Chemistry & biology 16, 548–556, doi:10.1016/j.chembiol.2009.04.009 (2009).

30 Eiamphungporn, W. & Helmann, J. D. The Bacillus subtilis sigma(M) regulon and its contribution to cell envelope stress responses. Mol Microbiol 67, 830–848, doi:10.1111/j.1365-2958.2007.06090.x (2008).

31 Tagami, K. et al. Expression of a small (p)ppGpp synthetase, YwaC, in the (p)ppGpp(0) mutant of Bacillus subtilis triggers YvyD-dependent dimerization of ribosome. MicrobiologyOpen 1, 115–134, doi:10.1002/mbo3.16 (2012).

32 Pedraza, J. M. & van Oudenaarden, A. Noise propagation in gene networks. Science 307, 1965–1969, doi:10.1126/science.1109090 (2005).

33 Milo, R., Jorgensen, P., Moran, U., Weber, G. & Springer, M. BioNumbers--the database of key numbers in molecular and cell biology. Nucleic Acids Res 38, D750–753, doi:10.1093/nar/gkp889 (2010).

34 Taniguchi, Y. et al. Quantifying E. coli proteome and transcriptome with single-molecule sensitivity in single cells. Science 329, 533–538, doi:10.1126/science.1188308 (2010).

35 Elowitz, M. B., Levine, A. J., Siggia, E. D. & Swain, P. S. Stochastic gene expression in a single cell. Science 297, 1183–1186, doi:10.1126/science.1070919 (2002).

36 Leveau, J. H. & Lindow, S. E. Predictive and interpretive simulation of green fluorescent protein expression in reporter bacteria. J Bacteriol 183, 6752–6762, doi:10.1128/JB.183.23.6752-6762.2001 (2001).

37 Radeck, J. et al. The Bacillus BioBrick Box: generation and evaluation of essential genetic building blocks for standardized work with Bacillus subtilis. J Biol Eng 7, 29, doi:10.1186/1754-1611-7-29 (2013).

38 Thackray, P. D. & Moir, A. SigM, an extracytoplasmic function sigma factor of Bacillus subtilis, is activated in response to cell wall antibiotics, ethanol, heat, acid, and superoxide stress. J Bacteriol 185, 3491–3498 (2003).

39 Salzberg, L. I. et al. The WalRK (YycFG) and sigma(I) RsgI regulators cooperate to control CwlO and LytE expression in exponentially growing and stressed Bacillus subtilis cells. Mol Microbiol 87, 180–195, doi:10.1111/mmi.12092 (2013).

40 Dubrac, S., Bisicchia, P., Devine, K. M. & Msadek, T. A matter of life and death: cell wall homeostasis and the WalKR (YycGF) essential signal transduction pathway. Mol Microbiol 70, 1307–1322 (2008).

41 Gaidenko, T. A., Kim, T. J. & Price, C. W. The PrpC serine-threonine phosphatase and PrkC kinase have opposing physiological roles in stationary-phase Bacillus subtilis cells. J Bacteriol 184, 6109–6114 (2002).

42 Libby, E. A., Goss, L. A. & Dworkin, J. The Eukaryotic-Like Ser/Thr Kinase PrkC Regulates the Essential WalRK Two-Component System in Bacillus subtilis. PLoS Genet 11, e1005275, doi:10.1371/journal.pgen.1005275 (2015).

43 Jones, D. L., Brewster, R. C. & Phillips, R. Promoter architecture dictates cell-to-cell variability in gene expression. Science 346, 1533–1536, doi:10.1126/science.1255301 (2014).

44 Sanchez, A., Garcia, H. G., Jones, D., Phillips, R. & Kondev, J. Effect of promoter architecture on the cell-to-cell variability in gene expression. PLoS Comput Biol 7, e1001100, doi:10.1371/journal.pcbi.1001100 (2011).

45 Shah, I. M., Laaberki, M. H., Popham, D. L. & Dworkin, J. A eukaryotic-like Ser/Thr kinase signals bacteria to exit dormancy in response to peptidoglycan fragments. Cell 135, 486–496 (2008).

46 Kriel, A. et al. GTP dysregulation in Bacillus subtilis cells lacking (p)ppGpp results in phenotypic amino acid auxotrophy and failure to adapt to nutrient downshift and regulate biosynthesis genes. J Bacteriol 196, 189–201, doi:10.1128/JB.00918-13 (2014).

47 Fisher, R. A., Gollan, B. & Helaine, S. Persistent bacterial infections and persister cells. Nat Rev Microbiol 15, 453–464, doi:10.1038/nrmicro.2017.42 (2017).

48 Hahn, J., Tanner, A. W., Carabetta, V. J., Cristea, I. M. & Dubnau, D. ComGA-RelA interaction and persistence in the Bacillus subtilis K-state. Mol Microbiol 97, 454–471, doi:10.1111/mmi.13040 (2015).

49 Germain, E., Castro-Roa, D., Zenkin, N. & Gerdes, K. Molecular mechanism of bacterial persistence by HipA. Mol Cell 52, 248–254, doi:10.1016/j.molcel.2013.08.045 (2013).

50 Levin-Reisman, I. et al. Antibiotic tolerance facilitates the evolution of resistance. Science 355, 826–830, doi:10.1126/science.aaj2191 (2017).

51 Ferrell, J. E., Jr. & Ha, S. H. Ultrasensitivity part II: multisite phosphorylation, stoichiometric inhibitors, and positive feedback. Trends Biochem Sci 39, 556–569, doi:10.1016/j.tibs.2014.09.003 (2014).

52 Nash, P. et al. Multisite phosphorylation of a CDK inhibitor sets a threshold for the onset of DNA replication. Nature 414, 514–521, doi:10.1038/35107009 (2001).

53 Zaytsev, A. V. et al. Multisite phosphorylation of the NDC80 complex gradually tunes its microtubule-binding affinity. Mol Biol Cell 26, 1829–1844, doi:10.1091/mbc.E14-11-1539 (2015).

54 Pufall, M. A. et al. Variable control of Ets-1 DNA binding by multiple phosphates in an unstructured region. Science 309, 142–145, doi:10.1126/science.1111915 (2005).

55 Shaffer, S. M. et al. Rare cell variability and drug-induced reprogramming as a mode of cancer drug resistance. Nature 546, 431–435, doi:10.1038/nature22794 (2017).

56 Kester, J. C. & Fortune, S. M. Persisters and beyond: mechanisms of phenotypic drug resistance and drug tolerance in bacteria. Crit Rev Biochem Mol Biol 49, 91–101, doi:10.3109/10409238.2013.869543 (2014).

57 Conlon, B. P. et al. Persister formation in Staphylococcus aureus is associated with ATP depletion. Nature microbiology 1, 16051, doi:10.1038/nmicrobiol.2016.51 (2016).

58 Gutierrez, A. et al. Understanding and Sensitizing Density-Dependent Persistence to Quinolone Antibiotics. Mol Cell 68, 1147–1154 e1143, doi:10.1016/j.molcel.2017.11.012 (2017).

59 Adams, K. N. et al. Drug tolerance in replicating mycobacteria mediated by a macrophage-induced efflux mechanism. Cell 145, 39–53, doi:10.1016/j.cell.2011.02.022 (2011).

60 Pu, Y. et al. Enhanced Efflux Activity Facilitates Drug Tolerance in Dormant Bacterial Cells. Mol Cell 62, 284–294, doi:10.1016/j.molcel.2016.03.035 (2016).

61 Geiger, T., Kastle, B., Gratani, F. L., Goerke, C. & Wolz, C. Two small (p)ppGpp synthases in Staphylococcus aureus mediate tolerance against cell envelope stress conditions. J Bacteriol 196, 894–902, doi:10.1128/JB.01201-13 (2014).

62 Wang, J. D., Sanders, G. M. & Grossman, A. D. Nutritional control of elongation of DNA replication by (p)ppGpp. Cell 128, 865–875, doi:10.1016/j.cell.2006.12.043 (2007).

63 Kriel, A. et al. Direct regulation of GTP homeostasis by (p)ppGpp: a critical component of viability and stress resistance. Mol Cell 48, 231–241, doi:10.1016/j.molcel.2012.08.009 (2012).

64 Krasny, L. & Gourse, R. L. An alternative strategy for bacterial ribosome synthesis: Bacillus subtilis rRNA transcription regulation. EMBO J 23, 4473–4483, doi:10.1038/sj.emboj.7600423 (2004).

